# Development of non-alcoholic steatohepatitis is associated with gut microbiota but not oxysterol synthesis

**DOI:** 10.1101/2022.12.02.518833

**Authors:** Jacqueline Wyss, Tina Raselli, Annika Wyss, Anja Telzerov, Gerhard Rogler, Niklas Krupka, Bahtiyar Yilmaz, Thomas SB Schmidt, Benjamin Misselwitz

**Affiliations:** Department of Visceral Surgery and Medicine, Inselspital, Bern University Hospital, University of Bern, Switzerland; Department of Gastroenterology and Hepatology, University Hospital Zurich and University of Zurich, Switzerland; Structural and Computational Biology Unit, European Molecular Biology Laboratory, Heidelberg, Germany; Maurice Müller Laboratories, Department for Biomedical Research, University of Bern, 3008, Bern, Switzerland

## Abstract

Liver steatosis is the most frequent liver disorder and its advanced stage, non-alcoholic steatohepatitis (NASH), will soon become the main reason for liver fibrosis and cirrhosis. The “multiple hits hypothesis” suggests that progression from simple steatosis to NASH is triggered by multiple factors including the gut microbiota composition. The Epstein Barr virus induced gene 2 (EBI2) is a receptor for the oxysterol 7a, 25-dihydroxycholesterol synthesized by the enzymes CH25H and CYP7B1. EBI2 and its ligand control activation of immune cells in secondary lymphoid organs and the gut. Here we show a concurrent study of the microbial dysregulation and perturbation of the EBI2 axis in a mice model of NASH.

We used mice with wildtype, or littermates with CH25H^-/-^, EBI2^-/-^, or CYP7B1^-/-^ genotypes fed with a high-fat diet (HFD) containing high amounts of fat, cholesterol, and fructose for 20 weeks to induce liver steatosis and NASH. Fecal and small intestinal microbiota samples were collected, and microbiota signatures were compared according to genotype and NASH disease state.

We found pronounced differences in microbiota composition of mice with HFD developing NASH compared to mice did not developing NASH. In mice with NASH, we identified significantly increased 33 taxa mainly belonging to the Clostridiales order and/ or the family, and significantly decreased 17 taxa. Using an Elastic Net algorithm, we suggest a microbiota signature that predicts NASH in animals with a HFD from the microbiota composition with moderate accuracy (area under the receiver operator characteristics curve=0.64). In contrast, no microbiota differences regarding the studied genotypes (wildtype vs knock-out CH25H^-/-^, EBI2^-/-^, or CYP7B1^-/-^) were observed.

In conclusion, our data confirm previous studies identifying the intestinal microbiota composition as a relevant marker for NASH pathogenesis. Further, no link of the EBI2 – oxysterol axis to the intestinal microbiota was detectable in the current study.

## Introduction

Non-alcoholic fatty liver disease (NAFLD) has a global prevalence of 25% (1), and currently constitutes the most frequent liver disorder. With global trends towards adaptation of a Western lifestyle worldwide, the number of people affected by NAFLD will rise even further. NAFLD comprises benign liver steatosis; however, it may progress to non-alcoholic steatohepatitis (NASH) (2) which can lead to liver fibrosis and liver cirrhosis with related complications including hepatocellular carcinoma. Since viral hepatitis has been substantially declining in the last decade due to the availability of effective antiviral treatments, NAFLD/ NASH will soon become the main reason for liver fibrosis and cirrhosis (3). Subsequently, NAFLD/ NASH will pose a significant clinical and economic burden on the healthcare systems (4,5).

The pathogenesis of NAFLD and NASH is critically linked to a metabolic state with overabundance of macronutrients including free fatty acids (FFA). NAFLD is characterized by synthesis and accumulation of triglycerides in the liver. Importantly, NAFLD is associated with other metabolic diseases such as obesity, type 2 diabetes mellitus and dyslipidemia, a constellation of conditions termed “metabolic syndrome” (6,7). NASH is characterized by a hepatic inflammatory reaction (8). However, only a subset of patients exhibiting similar metabolic comorbidities and dietary risk eventually develops NASH. The key steps mediating the progression of benign steatosis to NASH remain incompletely understood. A multiple hits hypothesis has been suggested in which additional adverse effects mediate hepatic inflammation (9). Main inflammatory triggers include the alteration of lipid metabolism, with changes in the generation of adipokines and cytokines and increased oxidative stress inside the liver. Further the gut microbiota has been suggested as one key factor for NASH pathogenesis (8,10).

A possible causal role of the gut microbiota in the generation of hepatic steatohepatitis has been suggested, as in mice NASH can be induced by transplantation of dysbiotic gut microbiota, whereas the transplantation of complete healthy gut microbiota can alleviate NASH susceptibility (11–13). The exact role of the gut microbiota in NASH pathogenesis is still unknown. Gut endotoxins, mainly bacterial lipopolysaccharides (LPS), have been shown to promote hepatic inflammation (14). Further possible hypotheses involve induction of a microbiota-induced gut barrier dysfunction by increasing epithelial permeability, shifts in bile acid composition influencing microbiota community structure, and bacterial metabolites, like ethanol, influencing hepatic metabolism (15). Increased permeability of the intestinal epithelium facilitates the transfer of bacterial lipopolysaccharides to the systemic circulation which can further promote proinflammatory effects (16).

Oxysterols are cholesterol metabolites, generated by enzymatic and non-enzymatic cholesterol oxidation (17). They constitute precursors for bile acid synthesis but have also been implicated in sterol homeostasis and immune regulation (18). One example of immune regulation via oxysterols is the EBI2-oxysterol axis.The Epstein Barr virus induced protein 2 (EBI2 – also known as GPR183) belongs to the family of G protein-coupled receptors (19). Two landmark papers have identified EBI2 as a high-affinity receptor for the oxysterol 7a, 25-dihydroxycholesterol (7a, 25-diHC), generated by the enzymes cytochrome P450 7B1 (CYP7B1) and cholesterol 25-hydroxylase (CH25H) (20,21). EBI2 and its ligand 7a, 25-diHC have been implicated in the positioning and the activation of B cells, T cells, and dendritic cells and the generation of an efficient antibody response (22–24). Further, the 7a, 25-diHC synthesizing enzyme CH25H has additional roles in inflammation and has been involved in intestinal fibrosis (25). Interestingly, no significant impact of CH25H, EBI2 and CYP7B1 knockout on murine NASH has been detectable in a previous study (26).

The role of oxysterols in NASH pathogenesis is not yet fully understood. An active role seems possible, as for instance hepatic oxysterols are elevated when animals are fed a diet high in free cholesterol (27). Further, oxysterols can have toxic effects on the liver by inducing mitochondrial dysfunction or by promoting inflammatory mediators like high mobility group box 1 protein (HMGB1) (28). An increase of 4b-OHC, 25-hydroxycholesterol (25-OHC), and 27-hydroxycholesterol (27-OHC) in NAFLD patients, (29), and an increase of 24-S-hydroxycholesterol (24(S)-OHC) and 7-hydroxycholesterol (7-OHC) derivates in NASH patients have been observed (26). Additionally, humans with a mutation of the cytochrome P450 7B1 (CYP7B1) enzyme are prone to develop liver cirrhosis and present with an accumulation of 24(S)-OHC, 25-OHC, and 27-OHC in the plasma (18). In murine models, a reduction of 27-OHC synthesis enhanced progression to NASH, whereas applications of subcutaneous 27-OHC had a protective effect (30).

The high degree of interconnection between metabolic state, oxysterol regulation, and microbiota, creates a challenge for disentangling individual effects. In the current study, we aimed to assess any concurrent relationship between players of the EBI2-oxysterol axis and microbiota composition in a murine-feeding model of NASH.

## Methods

### Murine feeding model

Approval by the local animal welfare authority (Tierschutzkommision Zürich, Zurich, Switzerland; registration number ZH 50/2013) was granted for the experimental protocol for animal testing. All animals used had a C57BL/6 background. Knockout animals for CH25H (Ch25h^-/-^), EBI2 (Ebi2^-/-^), and CYP7B1 (Cyp7b1^-/-^) were generated as previously described (26). The initial knockout animals were provided by Novartis Institutes (Ch25h^-/-^ and Ebi2^-/-^) or purchased from Jackson Laboratories (Cyp7b1^-/-^). Heterozygous mice were subsequently crossed with each other to obtain wild-type (Ch25h^+/+^, Ebi2^+/+^, or Cyp7b1^+/+^) and knockout (Ch25h^−/−^, Ebi2^−/−^, or Cyp7b1^−/−^) littermates. The mice used were eight-week-old littermates, housed under specific pathogen-free conditions in individually ventilated cages.

We induced NAFLD/NASH by a murine feeding model in C57BL/6 mice by supplying the animals for 20 weeks with a high-fat, high-cholesterol diet (HFD; ssniff Spezialdiäten GmbH) and a high-fructose corn syrup equivalent (55% fructose and 45% glucose, at a concentration of 42 g/l) in the drinking water as described previously (26). Control mice (standard diet, STD) were fed standard chow (Provimi Kliba) and water ad libitum. Wildtype and knockout animals were sacrificed and tested regarding clinical and histological hallmarks of NASH after 20 weeks (26). Further, microbiota samples from the small intestine and feces were collected. Liver samples were assessed by histology to score for the presence of steatosis, the occurrence of cellular hypertrophy or ballooning, and the presence of necroinflammation to diagnose NASH.

In total, 68 mice were investigated under HFD in the study and 42 fecal samples could be analyzed. From control animals under STD 54 fecal samples were included in the analysis.

**Table 1:** Overview of samples in regard to genotypes and NASH phenotype.

| **Genotype** | **NASH**<br>**(n=20)** | **no NASH**<br>**(n=22)** |
| --- | --- | --- |
| CH25H.WT | 3 | 1 |
| CH25H.KO | 3 | 3 |
| CYP7B1.WT | 5 | 3 |
| CYP7B1.KO | 9 | 6 |
| EBI2.WT | 2 | 4 |
| EBI2.KO | 0 | 3 |

### 16S rRNA amplicon sequencing

DNA was extracted using DNEasy PowerSoil kits (Qiagen, Hilden, Germany) as per the manufacturer’s instructions. Targeted amplification of the 16S rRNA V4 region (primer sequences F515 5’-GTGCCAGCMGCCGCGGTAA-3’ and R806 5’-GGACTACHVGGGTWTCTAAT-3’ (31), was performed in a two-step barcoded PCR protocol using the FailSafe PCR PreMix (Lucigen, WI, USA) according to the manufacturer’s instructions. PCR products were pooled, purified using size-selective SPRIselect magnetic beads (0.8 left-sized), and then sequenced at 2×250bp on an Illumina MiSeq (Illumina, San Diego, CA, USA) at the Genomics Core Facility, European Molecular Biology Laboratory, Heidelberg.

Raw 16S rRNA reads were trimmed, denoised, and filtered to remove chimeric PCR artefacts using DADA2 (32). The resulting Amplicon Sequence Variants (ASVs) were then clustered into Operational Taxonomic Units (OTUs) at 98% sequence similarity using an open-reference approach: reads were first mapped to the pre-clustered reference set of full-length 16S rRNA sequences at 98% similarity included with MAPseq v1.2.6 (33). Reads that did not confidently map were aligned to bacterial and archaeal secondary structure-aware SSU rRNA models using Infernal (34) and clustered into OTUs with 98% average linkage using hpc-clust (35), as described previously (36). The resulting OTU count tables were noise filtered by asserting that samples retained at least 400 reads and taxa were prevalent in at least 1% of samples; these filters removed 45% of OTUs as spurious, corresponding to 0.16% of total reads.

### Data analysis

Local sample diversities were calculated as OTU richness, exponential Shannon entropy and inverse Simpson index (corresponding to Hill diversities of order 0, 1, and 2 (37) as average values of 100 rarefaction iterations to 400 reads per sample. Between-sample community diversity was calculated as Bray-Curtis dissimilarity (38). Trends in community composition were quantified using ordination methods (Principal Coordinate Analysis, distance-based Redundancy Analysis) and tested using permutational multivariate analysis of variance (PERMANOVA (39) or ANOVA, as implemented in the R package *vegan* (40).

Machine learning models were built by randomly splitting data into test and training sets in 10 times repeated 10-fold cross-validation. For each fold, models were trained using the Elastic Net algorithm as implemented in the R package *siamcat* (41). Models were evaluated based on the average Area Under the Receiver Operating Characteristic curve (AUROC), averaged across validation folds. All codes used are deposited in the git repository at https://git.embl.de/tschmidt/ch25h-microbiome

## Results

### Impact of a high-fat diet on the intestinal microbiota

We used a high-fat diet (HFD) containing high amounts of fat and cholesterol with high-fructose corn syrup equivalent in the drinking water (42g/l) to induce NASH in mice. Animals were sacrificed at 20 weeks, and the degree of steatohepatitis was assessed in liver histology. As previously described, the HFD induced liver steatosis in most animals and NASH in approximately 50% of animals at 20 weeks, while the liver histology in controls remained normal, as described in detail before (26).

We analyzed the microbiota in fecal samples at 20 weeks by 16S sequencing. The HFD resulted in significant microbial changes in the mice. Taxa richness in stool samples of mice fed HFD was significantly lower than that of mice fed CHD (Fig 1A). This pattern persisted when richness was assessed by the exponential Shannon and the inversed Simpson diversity indices (S1 Fig). Diet further induced gross changes in the microbiota community structure. The ordination plot showed a clear distinction of the samples clustered together according to diet type (PERMANOVA R2=0.079, p≤10^-4; Fig 1B). Overall these results suggest that HFD diet impacts on the microbiota composition and leads to a reduction of alpha diversity.

**Fig 1:**
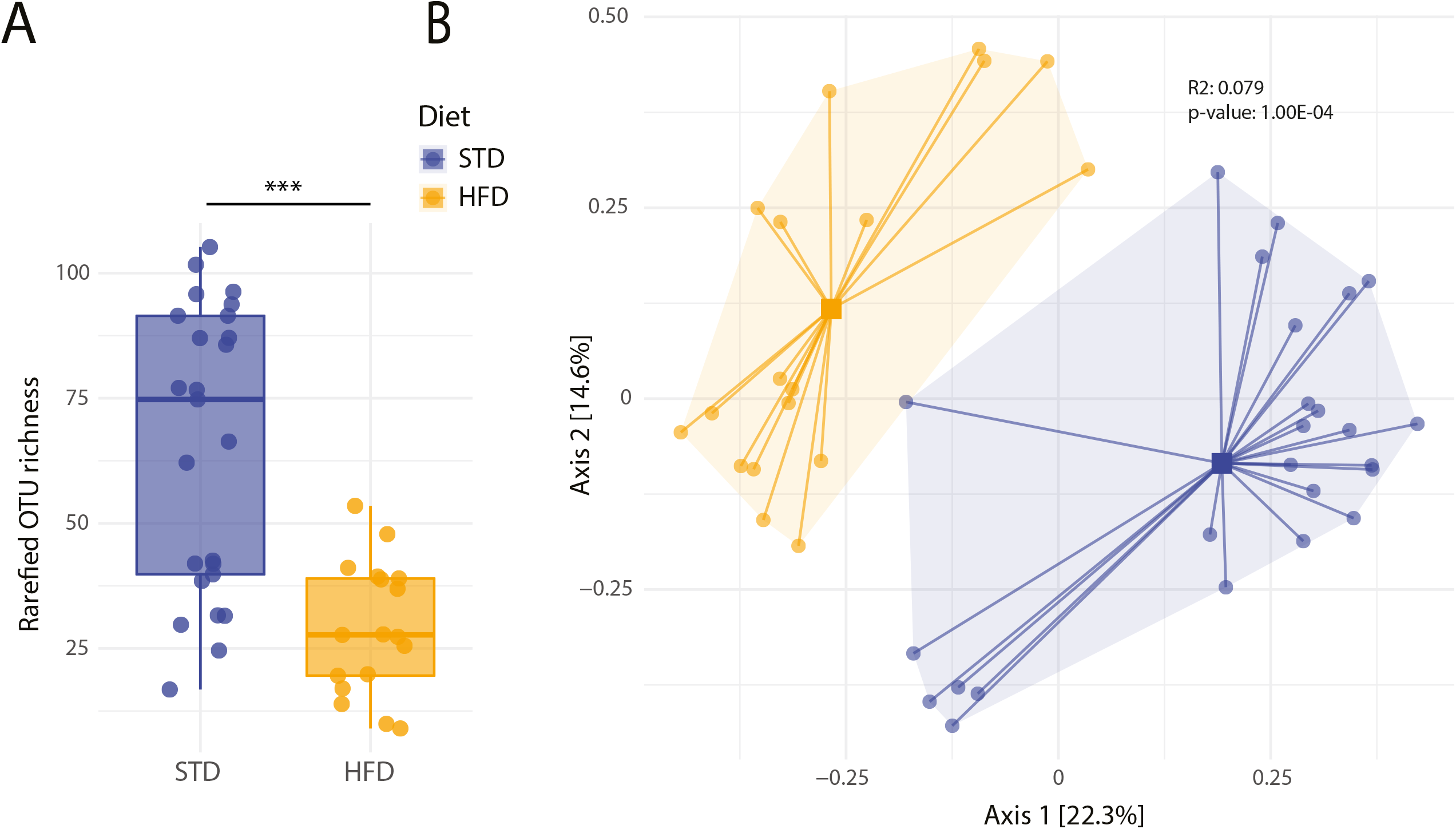
Comparison of species richness and beta-diversity of mice under standard diet and high-fat diet. A: Individual measurements of OTU richness after rarefaction to the lowest sequencing depth are depicted. Each dot represents a sample from a total of n= 43 mice. B: PCoA depiction of beta-diversity measured by Bray-Curtis dissimilarity with colors indicating samples belonging to the standard diet and the high-fat diet group.

### Microbiota profile in stool samples and the small intestine

To analyze the small intestinal microbiota, we isolated samples from the small intestine in mice with the HFD and STD controls. Species richness was higher in fecal samples than from samples collected from the small intestine; however, no difference in diversity was observed between the proximal, middle, and distal small intestines (Fig 2A, S2 Fig). In the ordination analysis, fecal samples and small intestinal samples were significantly separated, indicating a different microbiota composition (PERMANOVA R2=0.112, p≤10^-4). Further, samples from different sites of the small intestine clustered together, and a gradual shift from the proximal to the distal intestine was detected with samples from the distal small intestine being closest to fecal samples (Fig 2B).

**Fig 2:**
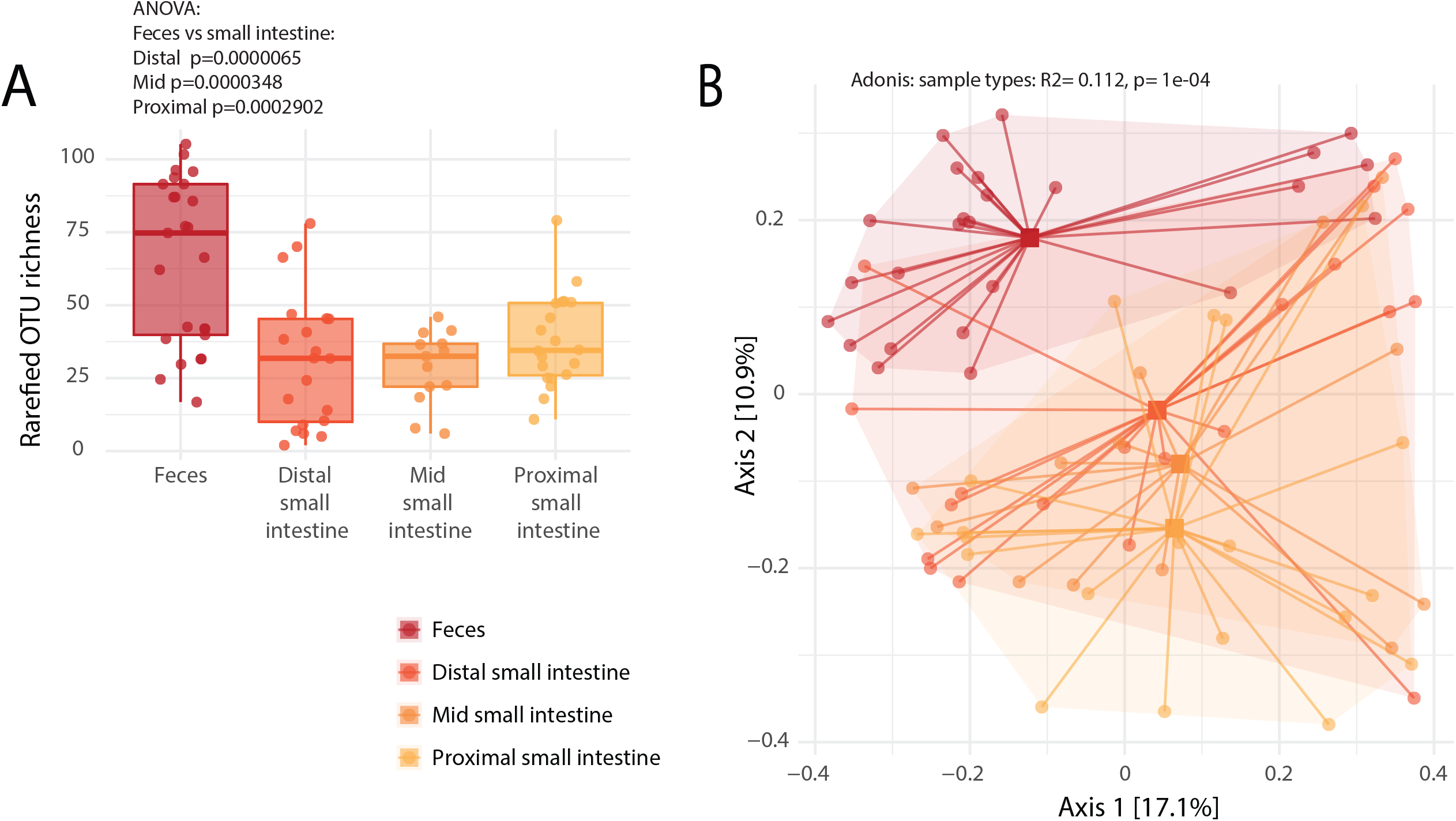
Comparison of species richness and beta-diversity of intestinal samples from different body sites. A: Individual measurements of OTU richness after rarefaction to the lowest sequencing depth are depicted. Each dot is representing a sample from a total of n= 77 mice. Plots are split according to sampling location. B: PCoA depiction of beta diversity measured by Bray-Curtis dissimilarity with colors indicating sampling location.

Similar to fecal samples, small intestine samples also tended to cluster according to feeding type and disease status in all small intestinal sampling locations (Fig 3A-C). However, the number of samples with sufficient recovered DNA from the HFD group was very low (n=27), thus limiting detailed comparisons. These findings indicate a systematic shift in the microbiota composition in regard to feeding type and the NASH status on top of an underlying gradient of the microbiota composition along the longitudinal axis of the intestinal tract.

**Fig 3:**
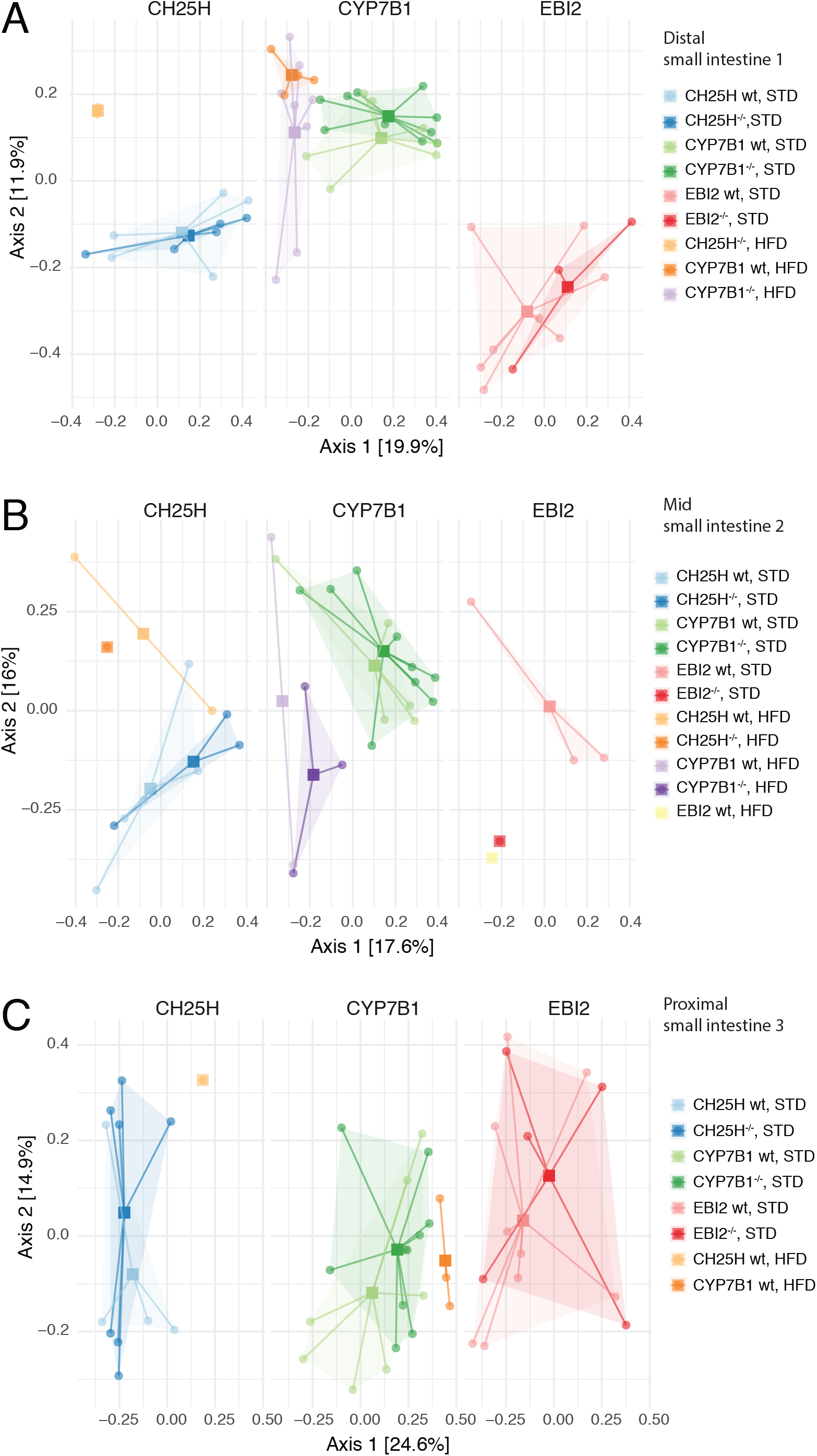
Comparison of beta-diversity according to diet type between intestinal samples stratified by genotypes. PCoA depiction of beta diversity measured by Bray-Curtis dissimilarity of genotypes and diet type. Samples are color-coded as indicated by the figure legend. Genotypes: wildtype (wt) and knockout. Diet: standard diet (STD) and high-fat diet (HFD). A: Distal small intestine. B: Mid small intestine. C: Proximal small intestine. The plot represents 14 HFD and 38 STD samples in the distal, 9 HFD and 26 STD samples in the mid, and 4 HFD and 42 STD samples in the proximal small intestine. Missing samples originate from technical inabilities to obtain successful sequencing results.

### Microbiota profile according to NASH disease state

Upon 20 weeks of HFD, approximately 50% of animals developed NASH. Interestingly, NASH was associated with a higher (although only borderline significant) microbiota diversity compared to animals with HFD without NASH (Figure 4A), indicating an association of the intestinal microbiota with liver inflammation beyond the HFD. The community analysis also indicated differences in microbiota community structures according to liver histology (PERMANOVA R2=0.108, p≤10^-4) even though we observed a pronounced overlap in the multidimensional scaling map (Figure 4B). Thus the change in alpha and beta diversity appears to be linked to NASH progression

**Fig 4:**
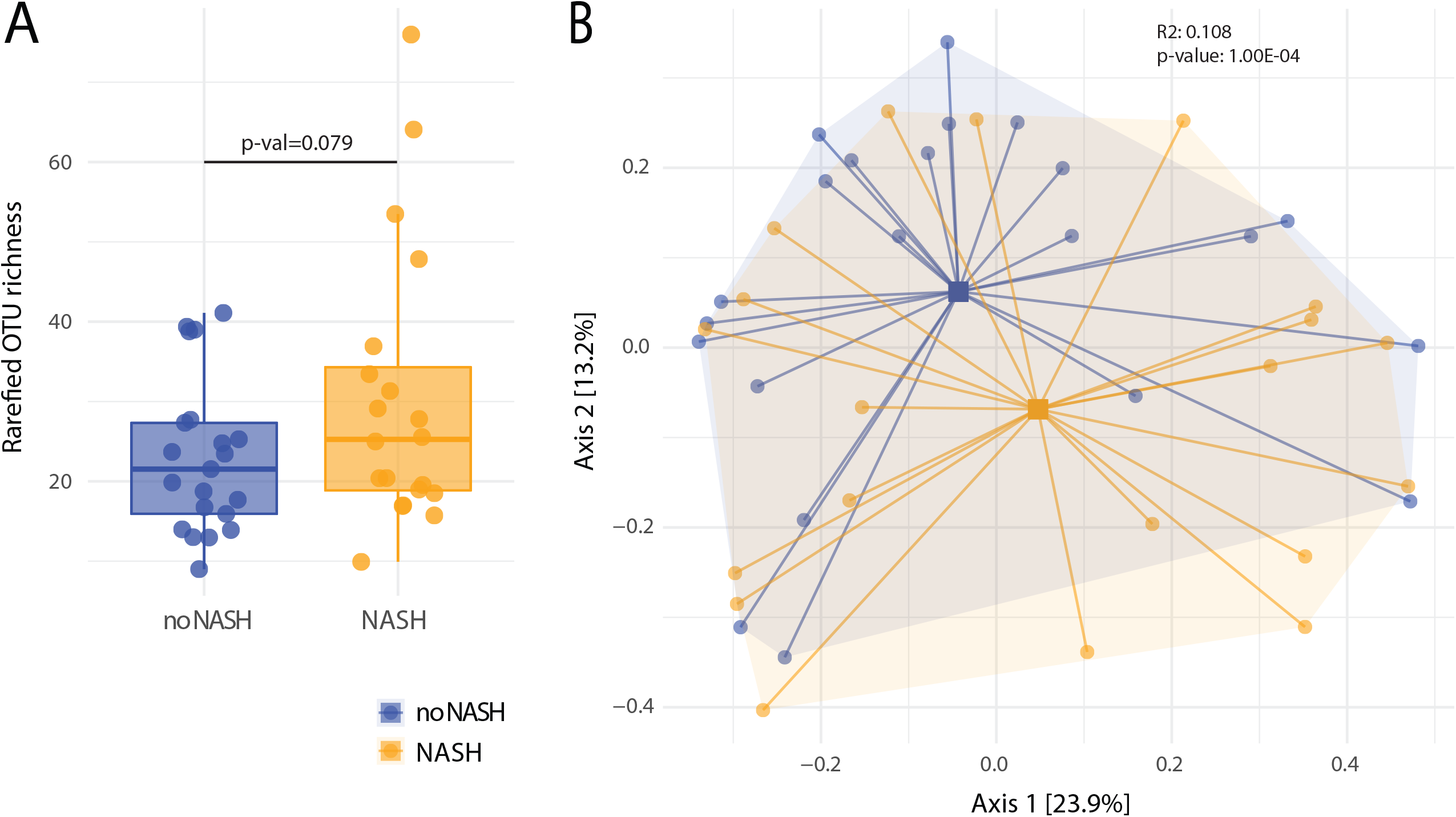
Comparison of species richness and beta-diversity of fecal samples according to NASH status. A: Individual measurements of OTU richness after rarefaction to the lowest sequencing depth are depicted. Each dot is representing a sample from a total of n= 42 mice. Plots are split according to disease status. B PCoA depiction of beta-diversity measured by Bray-Curtis dissimilarity with colors indicating NASH status of mice.

### Microbiota profile according to genotypes

Animals in our HFD trial comprised wildtype animals but also littermates with knockouts in the oxysterol receptor EBI2 or one of the 7a, 25-diHC synthesizing enzymes CH25H or CYP7B1. When the fecal samples were stratified by diet, NASH status and genotypes, microbiota clustered distinctly according to diet and NASH (Figure 5A-C) without apparent effects of the investigated genotypes wildtype, CH25H, EBI2, or CYP7B1. Similarly, samples from the distal, mid and proximal small intestine also showed no or only minimal deviation according to genotype (Supplementary Figure 3C-F).

**Fig 5:**
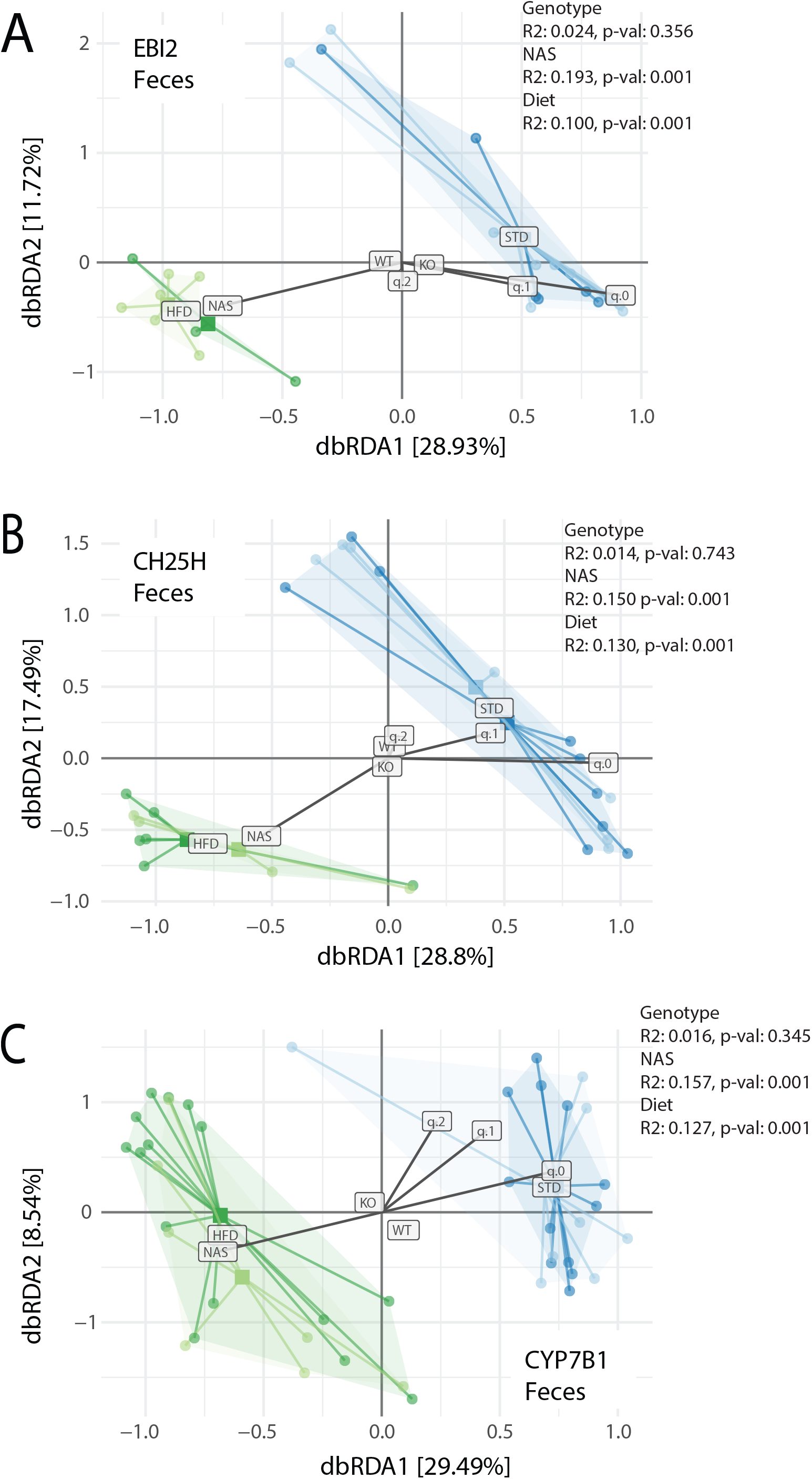
Distance-based redundancy analysis of fecal microbiota samples according to genotypes with diet, genotype, and NASH status as explanatory variables. Samples are colored according to feeding type (blue: standard diet, green: high-fat diet). A: EBI2^-/-^ (dark/light) versus wildtype. B: CH25H^-/-^ (dark/light) versus wildtype. C: CYP7B1^-/-^ (dark/light) versus wildtype. In these plots global centroids for genotype (’KO’ vs ‘WT’) and diet (’STD’ vs ‘HFD’) are shown; a larger distance of the centroids from ‘[0, 0]’ indicate a stronger effect on community composition. The (constrained) vectors of continuous variable effects are shown (for NAS, q.0, q.1, q.2); the direction of the vectors relative to each other indicates if effects are correlated (same direction) or anti-correlated (different direction). q.0, q.1, and q.2 are Hill alpha-diversities; q.0 is (rarefied) taxa richness, q.1 and q.2 are effective taxa numbers weighted by taxa abundances.

### Differential abundance analysis

We observed a significant increase of 33 OTUs and a decrease of 17 OTUs in animals with NASH compared to animals without NASH (Wilcoxon test, FDR-corrected p<0.05). Among the enriched OTUs, 17 belong to either the Clostridiales order and/ or the *Clostridiaceae* family, 3 were belonging to the Bacteroidales order, 2 belonging to the *Rikenellaceae* family, one was identified as *Akkermansia muciniphila*, 2 were belonging to *Ruminococcaceae* family, two were *Lactobacillus*, one belonged to the *Prevotellaceae* family, one was an *Eubacterium*, another belonged to the *Eubacteriaceae*, two to the *Oscillospiraceae* and one to the *Eggerthellaceae* family (Fig 6).

**Fig 6:**
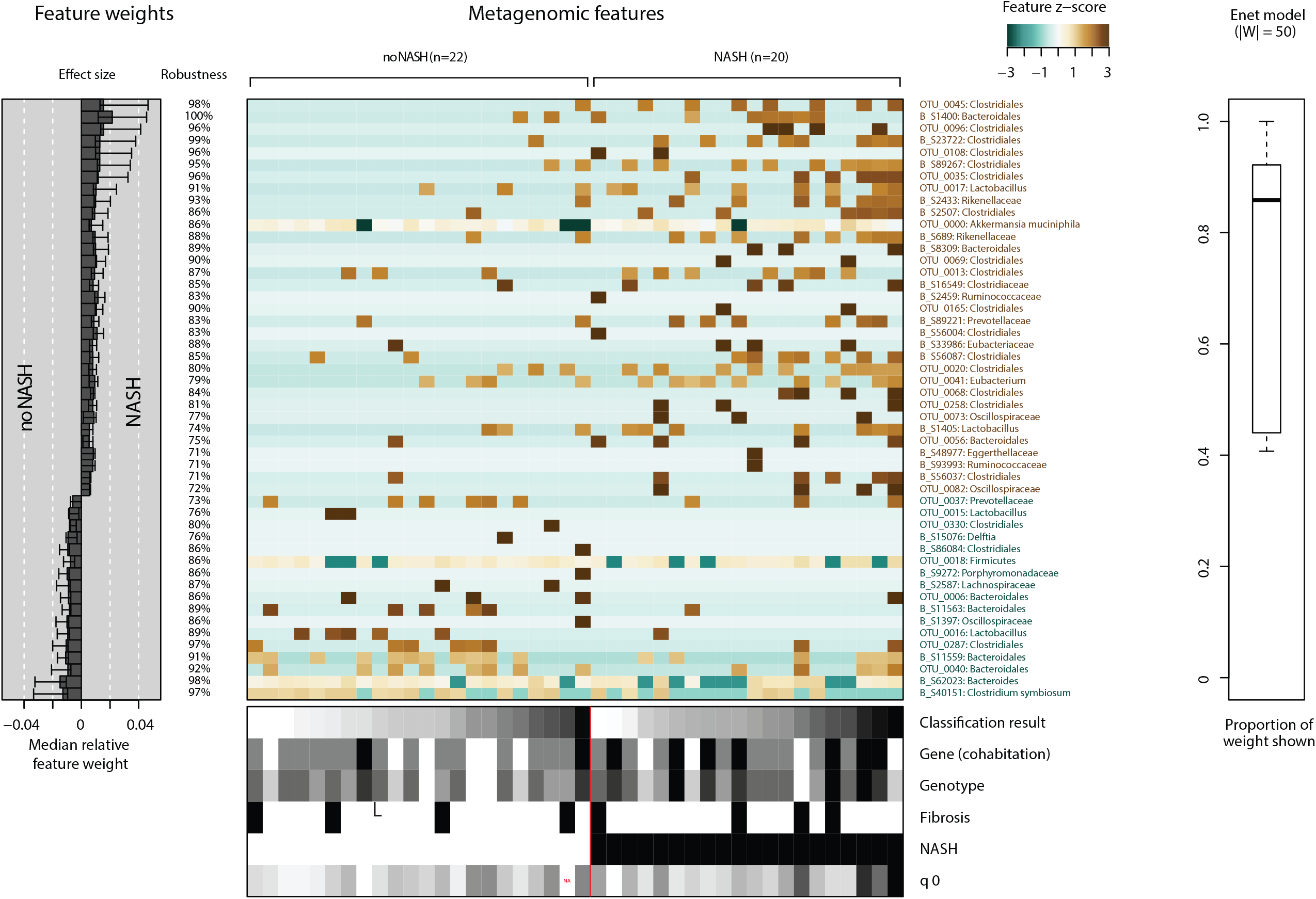
Identification of microbiota signatures according to NASH status after classification by SIMCAT. The importance of specific microbial taxa for the classification according to NASH status is shown. The weight of individual taxa used for classification is indicated together with individual effect size and robustness of classification. OTUs are named by their lowest taxonomic resolution and are colored according to their classification status (brown: NASH, green: no-NASH). The heatmap indicates taxa as features with sample abundance colored by the corresponding Z-scores. The panel at the bottom of the figures indicates the metadata available to the model (Classification: Result of Enet classification. Gene (cohabitation): Mice raised in the same cage. Genotype: wildtype vs knock-out CH25H^-/-^, EBI2^-/-^, or CYP7B1^-/-^. Fibrosis: Histology scoring of liver. NASH: Outcome measure of occurrence of NASH. q0: (rarefied) taxa richness).The panel with the title ENET model represents the weight statistic of the model (number of top features and their weight in the complete model).

Among the 17 OTUs depleted in NASH, 4 belong to *Bacteroidales*, one to *Bacteroides*, 2 to *Lactobacillus*, 3 to *Clostridiales*, 1 to *Clostridium symbiosum*, 1 to *Prevotellaceae*, one to *Delftia*, one belonging to Firmicutes, one to *Porphyromonadaceae*, one to *Lachnospiraceae* and one *Oscillospiraceae* (Fig 6).

To test how these univariate associations could be harnessed to predict NASH status, we next trained predictive models using the Elastic Net (ENET) algorithm using 10 times repeated 10-fold cross-validation (see Methods). The resulting models were moderately predictive of NASH at an AUROC of 0.64; interestingly, the algorithm exclusively picked microbiome features, as other features were homogenously distributed among the populations with and without NASH.

## Discussion

We provide evidence that fecal and intestinal microbiota composition in a murine feeding model of NASH differ between mice developing NASH and the ones that do not, according to liver histology upon feeding with a high fat diet. In mice with NASH, 33 OTUs were significantly enriched, mainly stemming from the Clostridiales order or the *Clostridiaceae* family, while 17 OTUs were significantly depleted. In contrast, the microbiota composition did not reveal strong differences between wildtype, CH25H^-/-^, EBI2^-/-^, or CYP7B1^-/-^ animals.

The most striking difference between the enriched OTUs comparing NASH and non-NASH mice in our study was the generally higher abundance of members of the Clostridiales order or the *Clostridiaceae* family in the NASH mice, while some members were also decreased. This might reflect a high baseline abundance of these taxa in the murine intestinal microbiota under a high-fat diet. It has to be noted that several members of this clade are metabolically active and might therefore benefit from the high influx of easily accessible metabolites (42,43). Previous microbiota signatures found in mice with NASH induced by several feeding models proved inconsistent so far. Similar to our results, a differential expression of some taxa with a high abundance of *Lactobacillus* and *Akkermansia* has been described, (44,45) while other groups reported differing findings (44–48).

The microbiota composition between animals fed with the HFD compared to the STD was markedly different. The lower species richness of the intestinal microbiota under a HFD is lower in accordance with previous studies (49). On a taxonomic level, microbiota signatures in mice provided with an HFD are consistently characterized by an increase of Firmicutes and a decrease of Bacteroidetes (50). On lower taxonomic levels, findings are more complex; however, in one study a swift and distinct increase of *Clostroidales, Verrucomicrobiales* and *Erysipelotrichiales* upon a high-fat diet challenge was a reproducible finding across multiple wildtype strains of mice (51). Animal studies have thus provided important insights into microbiota changes associated with NASH. The use of animal models permits standardization of the investigated subjects regarding sex, feeding patterns and in regard to metabolic comorbidities (especially type 2 diabetes) that are often difficult to control in human studies (52).

Our findings are in line with differences in human microbiota profiles in NAFLD and NASH compared to the composition in controls. Signatures differ from the ones observed in mice models. In human NASH, an enrichment of Proteobacteria has been consistently described (53–58). On the family level an increase of *Enterobacteriaceae* (53,56) and a decrease of *Rikenellaceae* (59) and *Ruminococcaceae* (53,54,60,61) has been observed. Several genera are consistently found to be increased (e.g. *Dorea* (54,59)) or decreased (*Faecalibacterium* (56), *Coprococcus* (56,59,62), *Anaerosporobacter* (62)). Furthermore, a differential abundance in *Lachnospiraceae* (53,54,59,62) has been commonly but less consistent described. To note, NASH shows some shared dysbiotic patterns similar to other metabolic diseases like obesity or type 2 diabetes mellitus (63) with decreased *Clostridium* levels (55,57,59,60) and increases in *Lactobacillaceae* including *Lactobacillus* (54,57,59,60,62) which in turn are shared with the profile detected in the current study.

The comparison of wildtype mice with CH25H^-/-^, EBI2^-/-^, or CYP7B1^-/-^ animals did not reveal relevant differences in their microbiota composition. While this argues against a relevant impact of the EBI2-oxysterol axis on the microbiota composition in NASH, other oxysterols might still be relevant, especially since bile acids and other products of liver metabolism can shape the gut microbiota (64). Bile acids are modified by the intestinal microbiota and are reabsorbed as secondary bile acids which have important immune regulatory functions (65–67). The composition and amount of bile acid secreted has been shown to be influenced by dysbiosis (68), and concurrent changes of bile acid composition and microbiota aberrations have been found in human NASH patients with hepatocellular carcinoma (60). The upregulation of bile acids can thus have a direct effect on microbiota composition, favoring strains with better adaption to higher bile acid concentrations (69). Therefore, complex indirect effects of changes in sterol synthesis affecting the gut microbiota have to be considered in studies addressing the intestinal microbiota and hepatic sterol metabolism.

Therapeutic efforts in liver steatosis and NASH are currently centered around lifestyle modification. While these are effective in reversing early disease stages, patient compliance is often lacking, limiting therapeutic success (70). Additional medical therapies are currently under investigation (9,55,71), potentially preventing and reversing NAFLD; however, no effective treatment for fibrosis is available yet. In end-stage liver disease, liver transplantation ultimately remains the only treatment option. Thus, understanding relevant factors of the gut microbiota influencing the progression of liver steatosis to NASH can potentially indicate new therapeutic targets.

Our study has several strengths and limitations. Strengths include the use of a relevant animal feeding model mimicking human NASH pathogenesis. Although generally NASH mice models do not show the full phenotype of human steatosis (especially hepatocyte ballooning) they are useful to investigate hepatic metabolic pathways (72). Notably, blinded histological analysis of mice livers in the current study confirmed histological hallmarks of NASH in our feeding model (26). Our study used littermates as controls, further reducing possible confounding factors.

An obvious limitation is the limited possibility of a direct translation of mice microbiota results to the human physiology. Further, human and murine oxysterol chemistry is complex, and our results only apply to EBI2 and its ligands and cannot be generalized to other oxysterols. Moreover, from many animals receiving a HFD, no sequencing results could be obtained in small intestinal samples, most likely for technical reasons. More generally, 16S amplicon sequencing based microbiome profiling provides a limited taxonomic resolution, although our analysis of both OTUs and ASVs exploited the available taxonomic profile information as far as currently possible.

In conclusion, differential expression of several intestinal microbiota taxa was detectable according to NASH disease state, confirming and expanding previous results. However, no impact of all tested genotypes (EBI2, CH25H, CYP7B1) on NASH could be detected, arguing against a relevant interaction of gut microbiota dysbiosis and the EBI – oxysterol axis in our murine NASH model.

## Abbreviations

NAFLD: non-alcoholic fatty liver disease
NASH: non-alcoholic steatohepatitis
FFA: free fatty acids
OTU: operational taxonomic unit
AUROC: area under the receiver operating characteristic curve
EBI2: Epstein Barr virus-induced gene 2 - also known as GPR183
CH25H: cholesterol 25-hydroxylase
CYP7B1: cytochrome P450 7B1
7a, 25-diHC: 7α,25-dihydroxycholesterol
25-OHC: 25-hydroxycholesterol
27-OHC: 27-hydroxycholesterol
24(S)-OHC: 24-S-hydroxycholesterol
7-OHC: 7-hydroxycholesterol
HFD: high fat diet
STD: standard diet

## Declarations

The authors would like to thank all technicians and animal carers for their commitment.

This work was supported by Swiss National Science Foundation Grant 32473B_156525 (B.M.), and a grant from the Hartmann-Müller Foundation (B.M.). The funding institutions had no role in the study design, analysis, interpretation of the data, or in writing the manuscript. *G*.*R. was supported by grants from AbbVie, Ardeypharm, MSD, Falk, Flamentera, Novartis, Roche, Tillots, UCB, and Zeller. B*.*M. has served on an advisory board for Takeda and BMS and has received traveling fees or speaking fees from Abbvie, MSD, Falk, iQONE, Gilead, and Novartis and has received unrestricted research grants from MSD and BMS unrelated to the current work*.

The authors declare that they have no competing interests. No third party has influenced any aspect of this study.

## Figure Legends

**S1 Fig: Comparison of samples between diet types according to alpha-diversity measures.** Comparison of samples from the standard diet and the high-fat diet by diversity indices (species richness, exponential Shannon index, inverse Simpson index) measured by the effective number of species.

**S2 Fig: Comparison of samples between body sites according to alpha-diversity measures.** Comparison of samples from intestinal sampling locations and feces by diversity indices (species richness, exponential Shannon index, inverse Simpson index) is shown, assessed by the effective number of species.

**S3 Fig: Distance-based redundancy analysis of small intestinal microbiota samples according to genotypes with diet, genotype, and NASH status as explanatory variables.** Samples are colored according to feeding type (blue: standard diet, green: high-fat diet). A: Distal small intestine, Ch25h^-/-^ (dark/light) versus wildtype (n=7 vs. 5). B: Distal small intestine, Cyp7b1^-/-^ (dark/light) versus wildtype (n=18 vs. 11). C: Mid small intestine, Ch25h^-/-^ (dark/light) versus wildtype (n=4 vs. 7). D: Mid small intestine, Cyp7b1^-/-^ (dark/light) versus wildtype (n=12 vs. 7). E: Proximal small intestine. Ch25h^-/-^ (dark/light) versus wildtype (n=7 vs. 5). F: Proximal small intestine, Cyp7b1^-/-^ (dark/light) versus wildtype (n=10 vs. 10). No analysis of Ebi2^-/-^ was feasible since only in a few small intestinal HFD samples from Ebi2^-/-^ animals (9 out of 33 HFD Ebi2^-/-^ animals) meaningful sequencing results were obtained.

