## Supplementary figures and images for "Development of non-alcoholic steatohepatitis is associated with gut microbiota but not oxysterol synthesis"

### Supplementary Figure S1

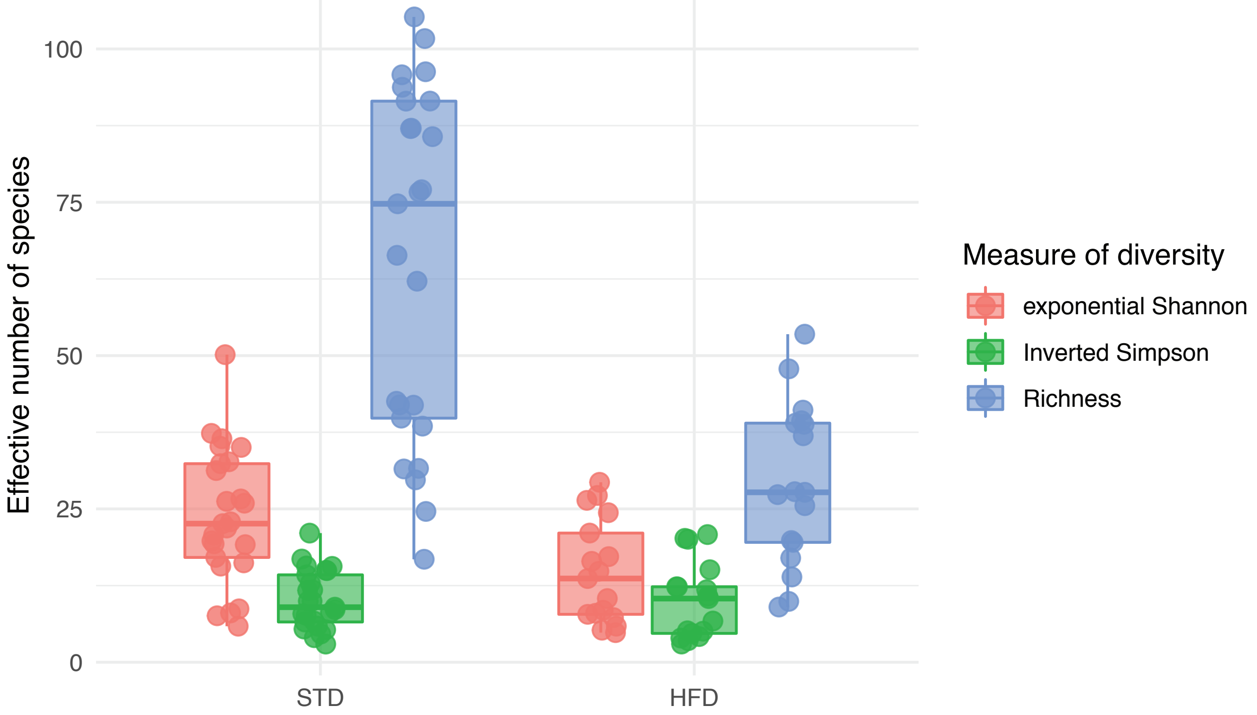

### Supplementary Figure S2

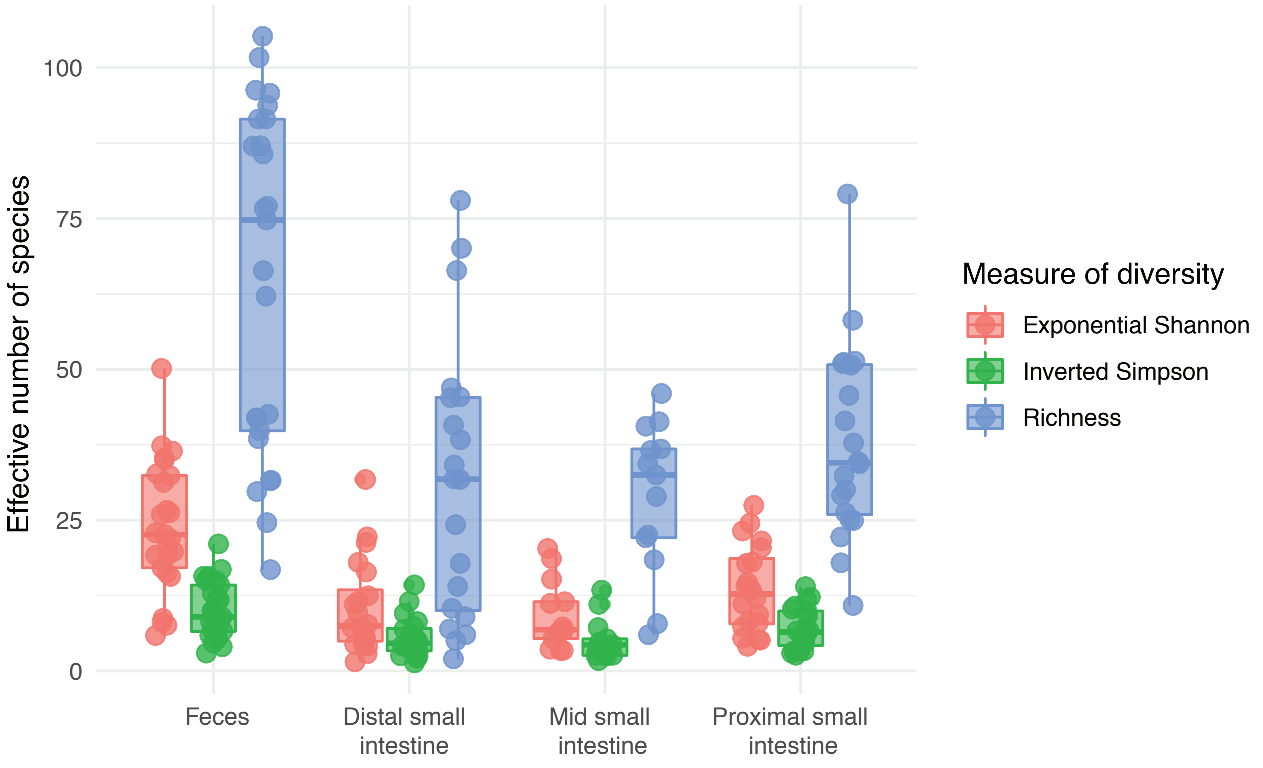

### Supplementary Figure S3

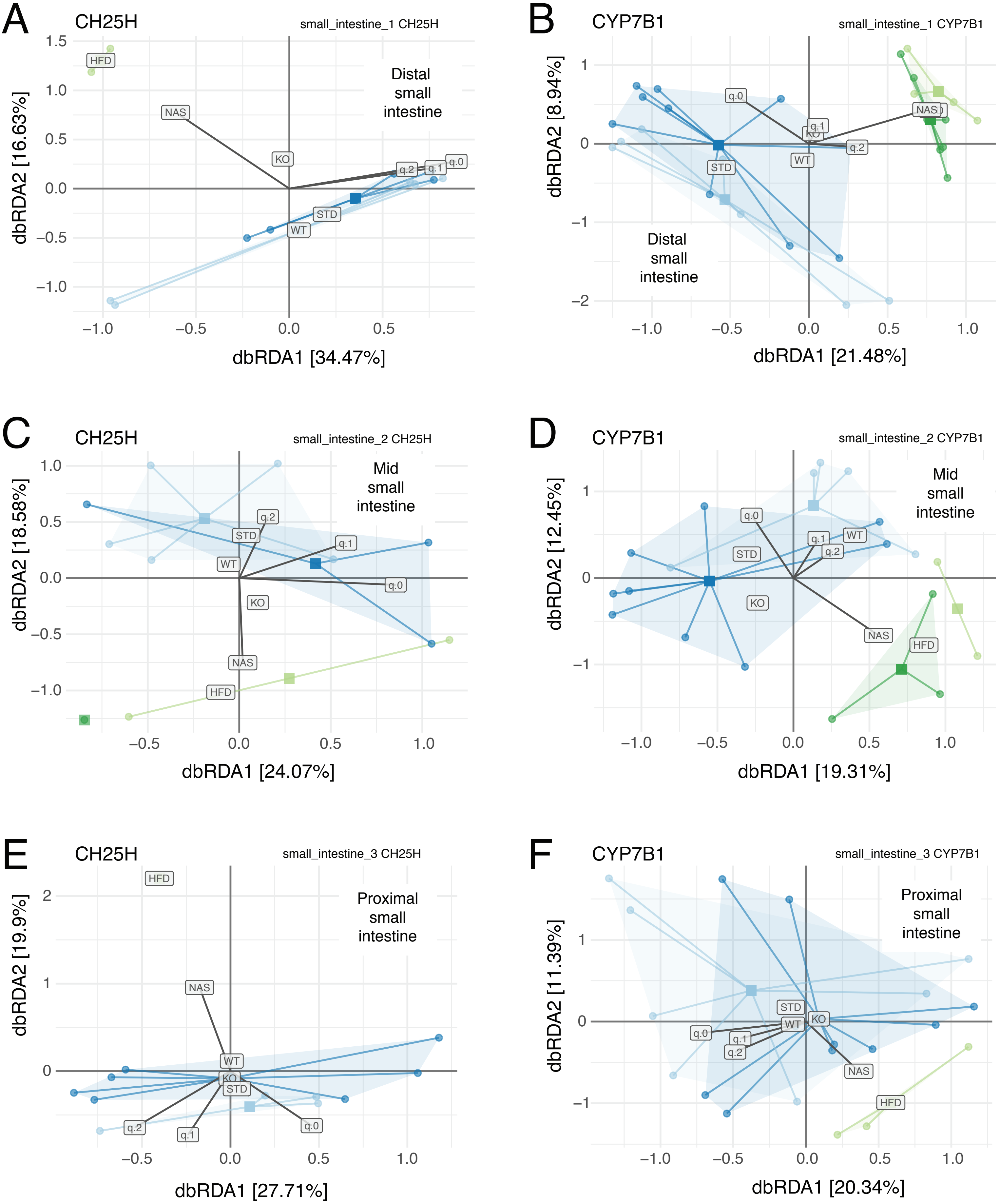
